# Novel genome characteristics contribute to the invasiveness of *Phragmites australis* (common reed)

**DOI:** 10.1101/2021.04.19.440155

**Authors:** Dong-Ha Oh, Kurt P. Kowalski, Quynh N. Quach, Chathura Wijesinghege, Philippa Tanford, Maheshi Dassanayake, Keith Clay

## Abstract

The rapid invasion of the non-native *Phragmites australis* (Poaceae, subfamily Arundinoideae) is a major threat to native ecosystems in North America. We describe a 1.14 Gbp reference genome for *P. australis* and compare invasive (ssp. *australis*) and native (ssp. *americanus*) genotypes collected across the Laurentian Great Lakes to deduce genomic bases driving its invasive success. We report novel genomic features including a lineage-specific whole genome duplication, followed by gene loss and preferential retention of genes associated with transcription factors and regulatory functions in the remaining duplicates. The comparative transcriptomic analyses revealed that genes associated with biotic stress and defense responses were expressed at a higher basal level in invasive genotypes, but the native genotypes showed a stronger induction of defense responses following fungal inoculation. The reference genome and transcriptomes, combined with previous ecological and environmental data, support the development of novel, genomics-assisted management approaches for invasive *Phragmites*.

## Introduction

Invasion of native ecosystems by non-native species is a worldwide problem damaging to ecosystems and economies. Invasive plants can negatively affect agricultural production ^1,2^ and displace native species through multiple mechanisms including enemy release ^3,4^, allelopathy ^5,6^, and novel traits ^7,8^. Thus, management and control of invasive plants can be a priority for conservation and agriculture.

*Phragmites australis* (Cav.) Trin. ex Steud.(Common Reed, Poaceae) is globally distributed (Fig. 1a) and provides multiple ecosystem services in its native range ^9–11^. A native subspecies (*P. australis* ssp. *americanus*) has been present in North American wetlands for thousands of years ^12^. However, the non-native, invasive subspecies (*P. australis* ssp. *australis*) ^13,14^ was introduced to North America from Europe around 1900 and has been aggressively disrupting and displacing native plant communities ^13,15^ and altering wildlife habitat and ecosystem properties ^16,17^. The invasive subspecies occurs throughout the contiguous United States (U.S.) and the entire Great Lakes basin ^13,18,19^ (Fig. 1b). It is one of the most problematic invasive plant species in wetland habitats in eastern North America, with hundreds of millions of dollars per year invested in control efforts ^20,21^. It is more robust than the native subspecies (Fig. 1d,e), with larger inflorescences, leaves, and height (Fig. 1f,g), but both subspecies reproduce by seed and clonally via rhizomes (Fig. 1h). A variety of mechanisms promoting *P. australis* invasions have been proposed, but effective control strategies are lacking ^22^. The Great Lakes Restoration Initiative identified invasive species (including *P. australis)* as one of its five most urgent issues ^23^. Invasive *P. australis* has also been recognized as the leading plant model for studying genetic mechanisms underlying plant invasions ^21,24^. *P. australis* therefore provides an excellent system to test genetic adaptations and control measures in plant invasions as both native and invasive populations coexist over a large geographic range (Fig. 1a,b).

**Fig. 1.**
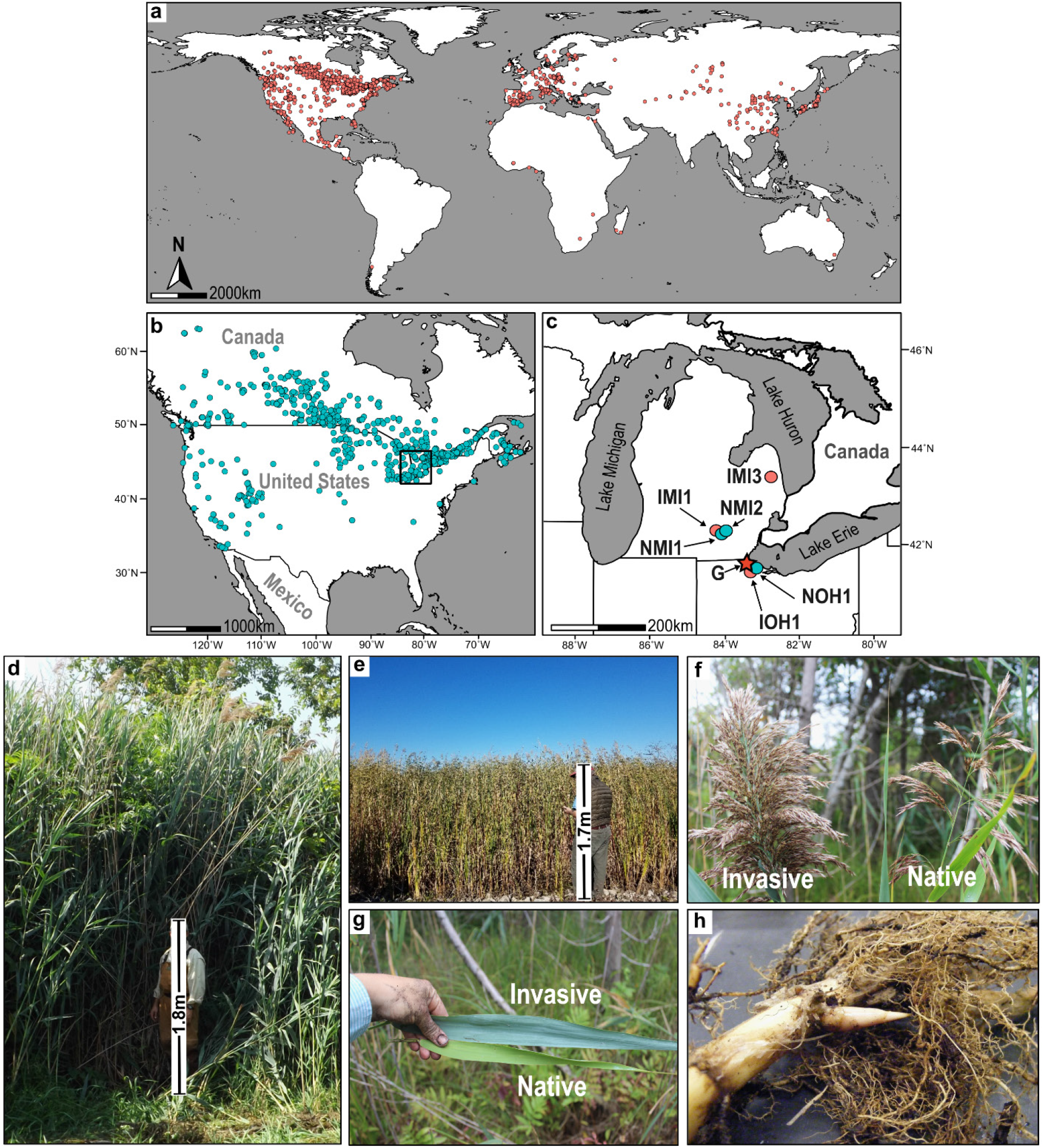
The invasive and native subspecies of common reed (*Phragmites australis*). **a,** Reported global distribution of *Phragmites australis* ssp. *australis* (“Invasive”) and **b,** Reported distribution of *P. australis* ssp. *americanus* (“Native”) in the U.S.A. and Canada based on the Global Biodiversity Information Facility (https://www.gbif.org/, data access September 2020). **c,** A magnified view of the box in panel **b** showing sample site locations in Michigan (MI) and Ohio (OH) in the U.S.A. Invasive (I) and Native (N) *Phragmites* plants were sampled from three Michigan sites (MI1, MI2, and MI3) and one Ohio site (OH1) in the Great Lakes region. Clonal fragments for the reference genome (G) were collected near the OH1 site. See Fig. 5a for more details. **d,** Invasive *P. australis* ssp. *australis* stand growing in a western Lake Erie coastal wetland. **e,** Native *P. australis* ssp. *americanus* stand located in a western Lake Erie wetland. **f,** Seed heads from the invasive and native *P. australis* growing in Michigan. **g,** Invasive and native *P. australis* leaves collected in Michigan. **h,** Rhizome and dense fibrous roots from an invasive *P. australis* plant growing in Michigan.

*P. australis* exhibits a range of ploidy levels from 2x-12x, with tetraploids reported as dominant in Europe and North America and octoploids dominant in Asia ^25,26^. *P. australis* genotypes, ploidy levels, and genome size have been assessed for traits that may favor invasiveness such as photosynthetic rate and nitrogen use efficiency, rhizome sizes, shoot emergence rates, and herbivory resistance ^27–29^. However, *P. australis* lacks a reference genome that can serve as a foundational resource to investigate genomic traits underlying plant invasions and to identify genetic targets for biocontrol. Further, *Phragmites* belongs to the grass subfamily Arundinoideae, which has been poorly explored at the genomic level compared to other grass subfamilies, even though some species are ecologically dominant or invasive on a global scale ^30^.

We report here the first reference genome for the invasive *Phragmites australis* ssp. *australis*, as well as comparative genomic and transcriptomic analyses that include invasive and native genotypes coexisting in the Great Lakes region of North America. Our results provide a key genomic resource for grasses and the subfamily Arundinoideae, novel insights into evolution in an underexplored grass clade, biological pathways correlated with invasiveness, and a genomic foundation for development of new management approaches.

## Results

### Assembly and annotation of the common reed *Phragmites australis*

We sequenced the genome of an invasive genotype of *Phragmites australis* ssp. *australis* collected in the U.S. Fish and Wildlife Service, Ottawa National Wildlife Refuge, Ohio (Fig. 1c, star-marked). We obtained ~42.19 Gbp of high confidence sequence data from leaf genomic DNA, representing 37-fold genome coverage, using PacBio SMRT sequencing technology. Assembled using Canu^31^ (version 1.4; see Methods), the reference genome provides 1.14 Gbp of 13,411 gap-free contigs with more than half of the assembled genome captured in 1,370 contigs (N50) larger than 194.6 kbp (L50). The largest contig is 3.22 Mbp (Table 1). Illumina short reads mapped to the primary assembly confirmed the genotype used as the reference being diploid based on single nucleotide polymorphisms that represented a non-reference allele frequency distribution with a peak at 0.5, as expected with a diploid genome (Supplementary Fig. 1). In total, 56.19% of the genome consist of repetitive sequences, with sequences derived from long terminal repeat (LTR) retrotransposons constituting 36.42% of the genome (Supplementary Table 1). Based on the repeat-masked genome sequence, we annotated 64,857 protein-coding gene models with a total length of 72.35 Mbp, which accounted for 6.35% of the genome (Table 1). The genomic location of these gene models and their best orthologs in rice and *Arabidopsis thaliana* are provided in Supplementary Dataset 1. BUSCO (Benchmarking Universal Single-Copy Orthologs) analysis ^32^ found 93.3% of single-copy gene models expected for land plants in the *Phragmites* reference genome (Supplementary Table 2).

**Table 1.** The *Phragmites australis* draft genome

|  |  |
| --- | --- |
| Assembly |  |
| Assembled genome size | 1,139,927,050 bps |
| Largest contig size | 3,219,705 bps |
| N50 contig length (L50) | 194,574 bps |
| Total number of contigs | 13,411 |
| Number of contigs > 1 Mbps | 67 |
| Number of contigs > N50 | 1,370 |
| Annotation |  |
| Number of protein-coding gene loci | 64,857 |
| Total length of predicted ORFs | 72,347,598 bps |
| Longest ORF | 14,790 bps |
| Median length of ORFs | 927 bps |
| Proportion of ORFs in the genome | 6.35 % |
| Proportion of repeats in the genome | 56.19 % |
| SINEs | 0.14 % |
| LINEs | 1.74 % |
| LTR elements | 36.42 % |
| DNA elements | 11.43 % |
| Unclassified repeats | 6.46 % |

We constructed a species tree using 6,404 gene models representing 782 gene families from *P. australis* and 13 other published grass genomes (PLAZA monocot database v. 4.5) ^33^, with pineapple (*Ananas comosus,* Bromeliaceae) as the outgroup. *P. australis*, representing the first genome from the Arundinoideae subfamily, was placed sister to subfamily Chloridoideae, consistent with the PACMAD clade species tree (Fig. 2) ^34,35^.

**Fig. 2.**
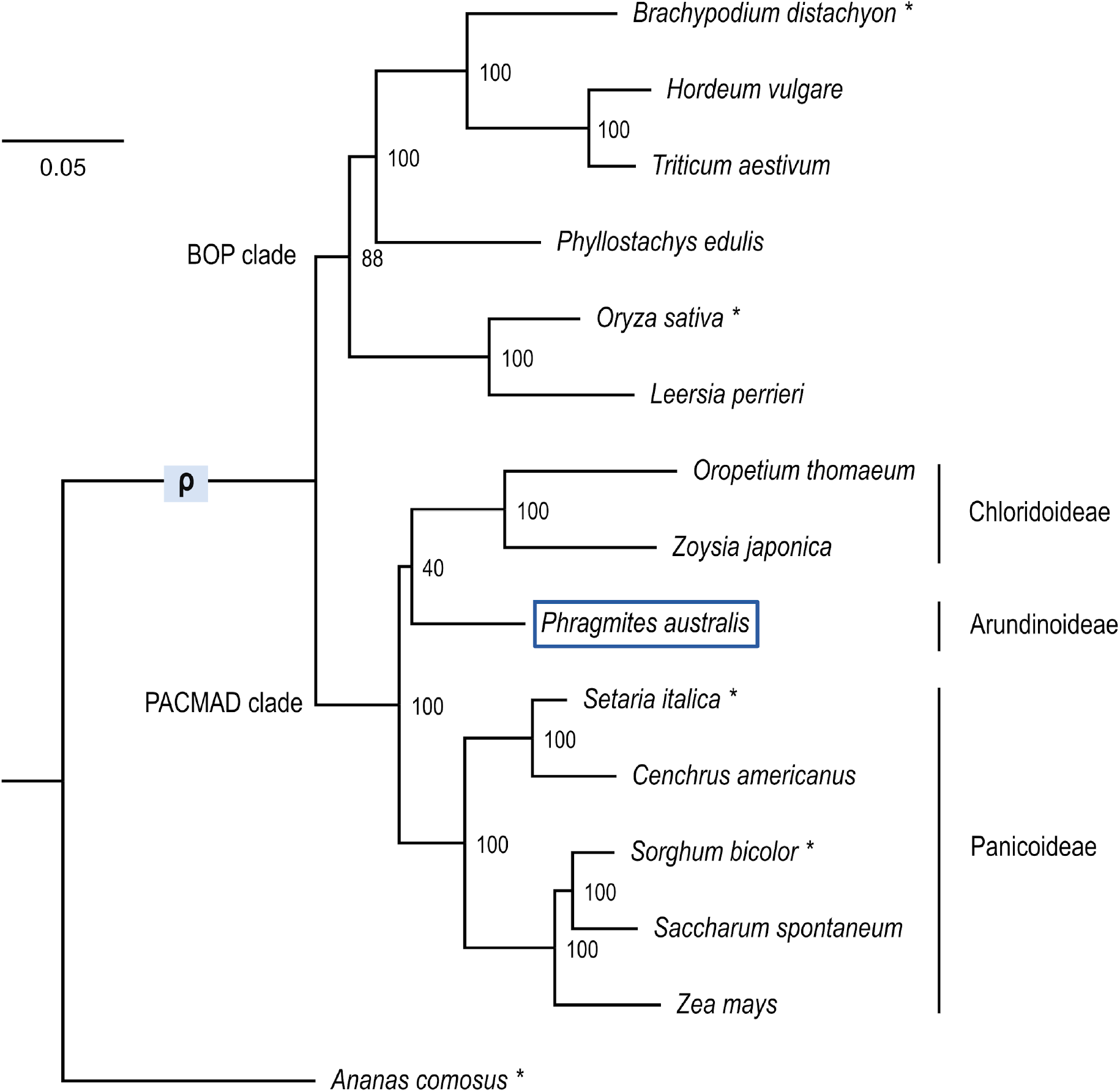
Phylogenetic position of *Phragmites australis* in the under-investigated Arundinoideae subfamily, based on the draft genome. For the *P. australis* draft genome and 14 monocot genomes publicly available in Monocots PLAZA (v. 4.5) ^33^, a maximum likelihood species tree was constructed based on 26,878 amino acid alignments from 782 ortholog groups, selected based on the criteria that at least seven species among the set had a single ortholog. All sites that included gaps in more than 20% of taxa were excluded. The number in each branch shows percent support from 1000 bootstrap replicates. The branch with the ρ genome duplication ^36^ is marked, and species with asterisks were used for comparative analyses.

### Signatures of a recent whole genome duplication in the *P. australis* genome

We found signatures of a previously unreported whole genome duplication in the *P. australis* genome that occurred after its divergence from the subfamily Panicoideae (Fig. 3). We compared the *P. australis* genome with five representative monocot reference genomes, including pineapple (*A. comosus*) and four grass species without a genome duplication more recent than the ρ duplication ^36^ (Fig. 2, marked with asterisks). The *P. australis* genome presented 36.7% BUSCO genes as duplicated, while only 1-5% BUSCOs were duplicated in the five comparator genomes (Fig. 3a and Supplementary Table 2).

**Fig. 3.**
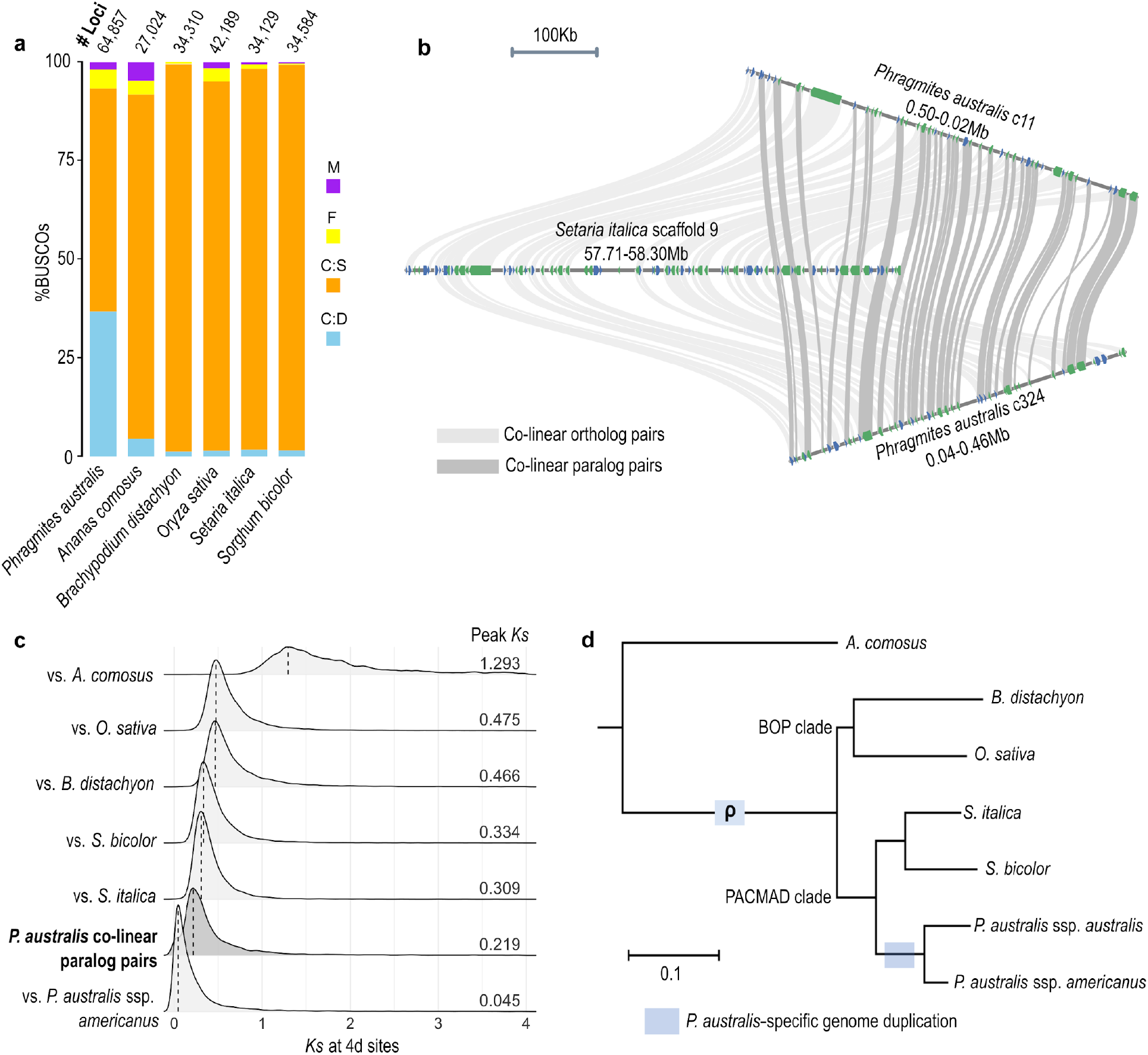
Signatures of *Phragmites australis*-specific whole genome duplication (WGD). **a,** Percentages of complete duplicated (C:D), complete single-copy (C:S), fragmented (F), and missing (M) orthologs among 1,375 Benchmarking Universal Single-Copy Orthologues (BUSCOs) and the number of protein-coding gene loci (# Loci), in the genomes of *P. australis* and other monocot species. **b,** An example microsynteny between a 500-Kb *Setaria italica* genomic block and two duplicated *P. australis* genomic blocks. Ribbons connect co-linear ortholog (light gray) and paralog (dark gray) pairs identified by MCscan^58^ as described in Methods. **c,** Synonymous substitution rates (*Ks*) at 4-fold degenerate (4d) sites were estimated for co-linear ortholog pairs between *P. australis* and *A. comosus* (total 2,720 ortholog pairs), *B. distachyon* (10,559 pairs), *O. sativa* (11,176 pairs), *S. italica* (12,878 pairs), and *S. bicolor* (12,322 pairs), respectively, as well as 6,600 co-linear paralog pairs detected within the *P. australis* genome. For comparison with the native *P. australis* ssp. *americanus*, we used 11,445 reciprocal best homolog pairs between *P. australis* reference genome and a *de novo* transcriptome assembly from *P. australis* ssp. *americanus*. Probability distributions and peak values of *Ks* are shown. **d,** A maximum-likelihood species tree shows the branches associated with the *P. australis*-specific WGD, as well as the ρ WGD event shared among grasses ^36^.

A genome-wide alignment between *Setaria italica* and *P. australis* found that 50.2% of *S. italica* gene models are represented twice in the *P. australis* genome as co-linear orthologs in synteny blocks (Fig. 3a, and Supplementary Figure 2a). Similar patterns were observed in comparisons between *P. australis* and the other four monocot genomes (Supplementary Figure 2a).

A genome-wide self-alignment using SynMap identified 14,005 gene loci, comprising 21.6% of all *P. australis* protein-coding genes, that were organized into 1,501 paralogous synteny blocks consisted of at least five co-linear paralog pairs. Synteny blocks were widespread across the *P. australis* genome and found in 66.8% of all contigs with ten or more protein-coding gene loci (Supplementary Fig. 2b and c), further supportive of whole genome duplication (WGD) events.

To assess the timing of genome duplications that resulted in the observed paralogous synteny blocks, we used neutral evolutionary substitutions calculated for 4-fold degenerate (4D) sites in codons that allows positioning of timing of duplicated events within a clade. We plotted the distribution of synonymous substitution rate (*Ks*) at 4D sites for *P. australis* co-linear paralog pairs and for co-linear ortholog pairs between *P. australis* and comparator species. (Fig. 3c). The peak *Ks* for *P. australis* co-linear paralog pairs (Fig. 3c, dark gray) was smaller than those observed for any pairwise comparisons between species, while larger than the value found between *P. australis* ssp. *australis* and native *P. australis* ssp. *americanus* (Fig. 3c, bottom row). We also found a small proportion of synteny blocks within the *P. australis* genome that likely represent the ρ duplication at the root of the Poaceae (Supplementary Fig. 3). Our analysis suggests that the majority of co-linear paralogs in synteny blocks are derived from a single WGD event that occurred after the divergence of *Phragmites* from the Panicoideae, represented here by *S. italica* and *S. bicolor,* but before the divergence between the subspecies *australis* and americanus (Fig. 3d).

### Biased retention of transcription factors and signaling-related genes following the lineage-specific genome duplication in *P. australis*

While the synteny blocks are widespread across the *P. australis* genome, the relatively short length of each synteny block (Supplementary Fig. 2c) and the fact that only 21.6% of duplicated genes were retained in synteny blocks indicates a substantial fractionation following genome duplication, likely due to the selective retention of duplicates that enhance plant fitness and loss of redundant or detrimental duplicates. We therefore compared the functions of duplicated *P. australis* genes relative to those genes with conserved copy numbers based on ortholog groups including other grass species (Fig. 4).

**Fig. 4.**
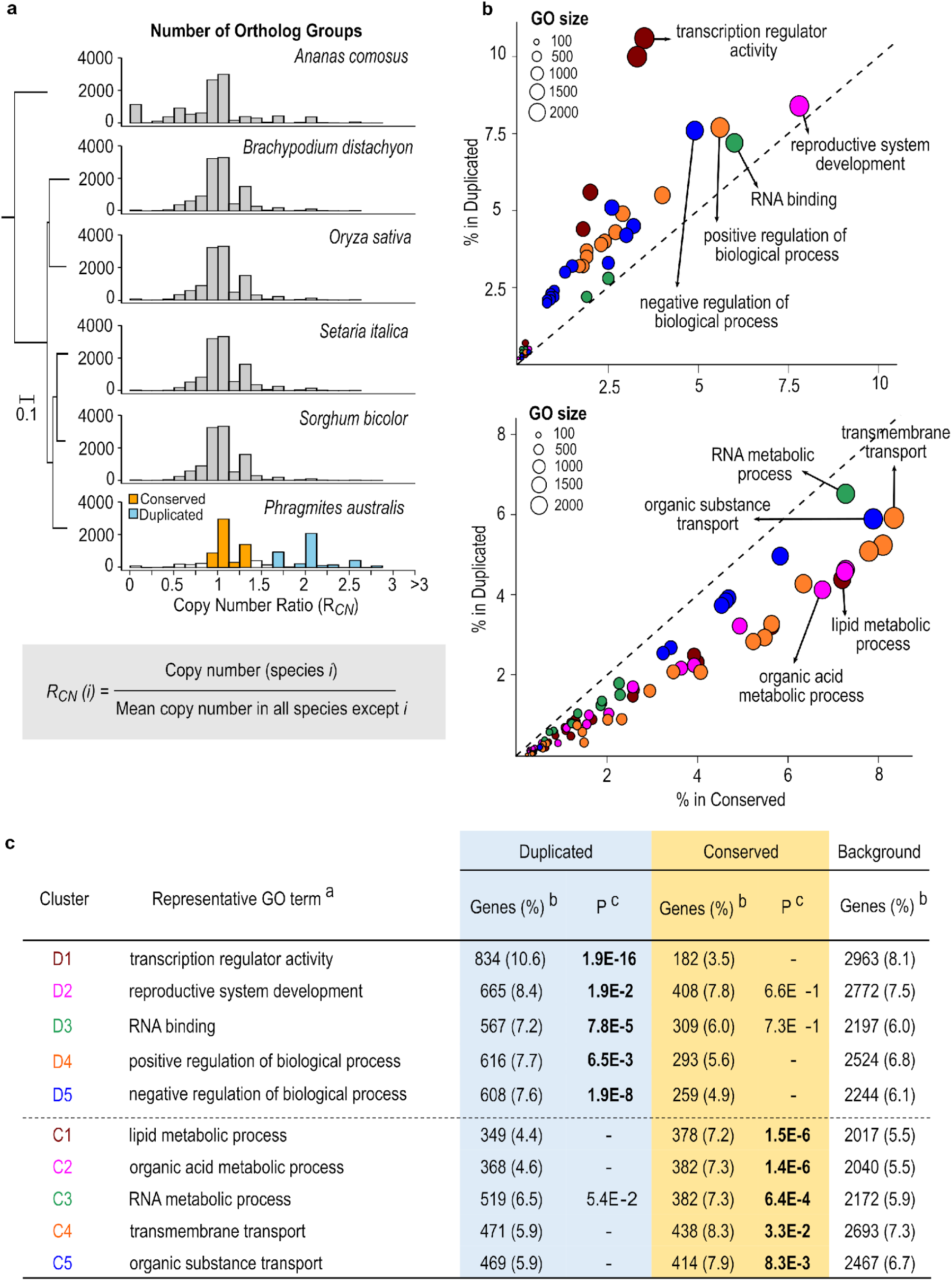
Functions enriched among *P. australis* genes that remained duplicated after the WGD event. **a,** Comparison of ortholog copy numbers between *P. australis* and five other monocot species. For each ortholog group identified by OrthoFinder as described in Methods, Copy Number Ratio (*R*_*CN*_) was calculated for each species by dividing the ortholog number in the species with the mean ortholog copy number in the other five species. Histograms show a shift towards increased *R_CN_* uniquely in *P. australis*. We identified “Conserved” (orange) and “Duplicated” (sky-blue) groups among *P. australis* genes whose copy numbers remain unchanged and uniquely increased, respectively, compared to other monocot species. **b-c,** GO terms enriched in either Duplicated or Conserved *P. australis* gene groups. The proportion of genes annotated with each GO term in Duplicated and Conserved groups was plotted as circles in (**b**). GO terms at least 80% overlapping with a bigger GO term are clustered and the largest five GO clusters enriched in either group were shown with the same color in (**b**) and (**c**). ^a^ The largest GO term in each GO cluster; MF, Molecular Function; BP, Biological Process. ^b^ Percentages are to the total number of genes with a GO annotation in each group. ^c^ P-values of enrichment compared to the Background, after Benjamini-Hochberg correction for multiple testing. Values less than 0.05 are in bold.

For all ortholog groups (OGs) detected among *P. australis* and the five comparator monocot species (Methods and Supplementary Dataset 2), we calculated the ortholog copy number ratio (*R_CN_*) for each species by dividing the copy number in the species with the average of the other five species (Fig. 4a and Supplementary Table 3). All species except *Phragmites* showed a peak at *R_CN_* ≈ 1. By contrast, the *R_CN_* distribution in *P. australis* showed two peaks, one with *R_CN_* at 1 (Fig. 4a, “Conserved”) and a second peak at *R_CN_* ≈ 2 (Fig. 4a, “Duplicated”), indicating that a substantial number of *P. australis* OGs have their copy numbers doubled compared to other species (See Methods for details). We searched for enriched gene functions in the Duplicated group (11,002 *P. australis* genes in 4,113 OGs) and in the Conserved group (6,981 *P. australis* genes in 5,600 OGs). Fig. 4b, c present the five largest functional clusters, detected using GOMCL ^37^, considering GO terms significantly enriched exclusively in either Conserved or Duplicated groups. The largest clusters included 72% and 64% of all *Phragmites* genes represented by 123 and 146 GO terms enriched in the Duplicated and the Conserved groups, respectively (Supplementary Dataset 3). Functions enriched exclusively in the Duplicated group were largely related to regulation of gene expression (Fig. 4b, c). For example, “transcription regulator activity” is the most enriched functional cluster among duplicated genes (Fig. 4b, c, cluster D1), representing 10.6% of all genes in the Duplicated group with a GO annotation, compared to only 3.5% and 8.1% in the Conserved group and all genes (used as the background for the enrichment analysis), respectively (Supplementary Dataset 3). By contrast, the Conserved group was enriched in functions associated with primary metabolic processes and transport (C1-5 in Fig 4b, c and Supplementary Dataset 3). Selective gene retention after WGD events are often linked to adaptive traits or traits leading to lineage diversification. The biased retention of transcription factors and regulatory processes in *P. australis* spp. *australis* following its most recent WGD event is indicative of greater genomic plasticity conducive to invasive lifestyles ^38^.

### Divergence in basal transcriptome profiles between invasive and native *P. australis* subspecies

To gain further insight into genetic factors promoting invasiveness, we compared transcriptomes of invasive and native *Phragmites* subspecies collected from the Great Lakes region (Fig. 1c, 5a). We generated eight replicates of RNA-seq samples from leaf and rhizome tissues, respectively (see Methods), followed by independent *de novo* transcriptome assembly of the six genotypes (Supplementary Table 4). The maximum likelihood tree based on concatenated alignments of 3,464 homolog groups separated the six genotypes into invasive and native subspecies as expected, and the three invasive genotypes were placed together with the invasive genotype used as the reference genome (Fig. 5b).

**Fig. 5.**
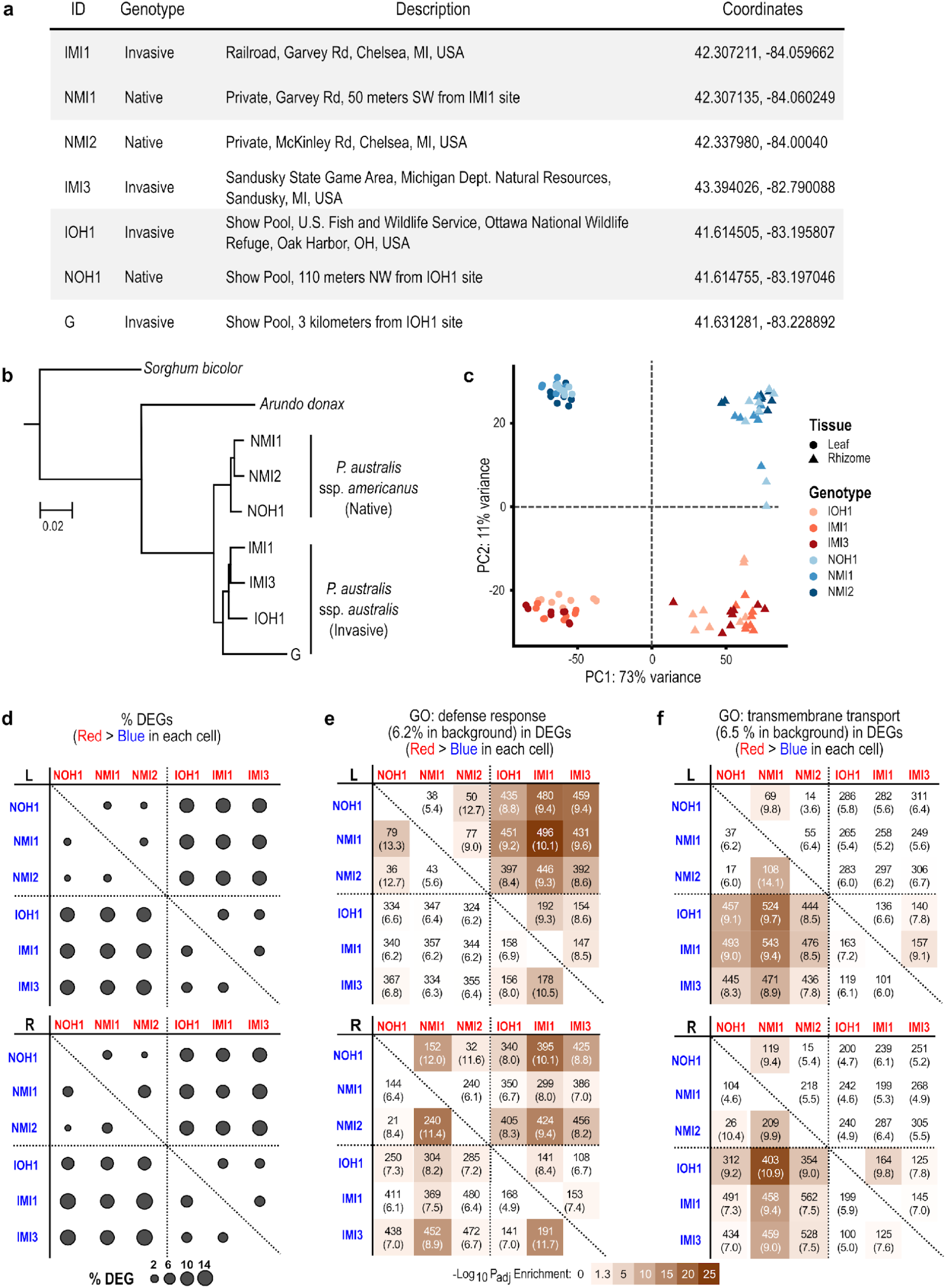
Comparison of transcriptomes between invasive and native *P. australis* genotypes. **a,** Details on genotype collections of invasive (I) and native (N) subspecies from locations marked on Fig. 1c. **b,** A maximum likelihood tree based on deduced protein sequences of 5,394 homologous gene groups. Protein sequences were deduced from transcriptomes *de novo* assembled using RNA-seq reads derived from the six *P. australis* genotypes shown in (**a**) as well as the reference gene models (marked with “G”). Publicly available transcriptome and genome sequences were used for *Arundo donax* ^69^ and *Sorghum bicolor*, respectively. All branches were 100% supported by 100 bootstrap tests. **c,** RNA-seq reads from leaf and rhizome tissues of the six genotypes were aligned to the *P. australis* reference genome. Principal Component Analysis separated different tissues and genotypes. **d,** Differentially Expressed Genes (DEGs) showing significant (adjusted p-value < 0.05 and fold-difference ≥2) changes in basal-level expression in leaf (L) and rhizome (R) were identified between all pairs of genotypes. In the diagonal plot, the circle in each cell represents the proportion of DEGs among 64,857 *P. australis* reference gene models in which the genotype in the column (red) shows higher basal-level expression than the clone in the row (blue). **e-f,** GO terms “defense response” (**e**) and “transmembrane transport” (**f**) showed enrichment among DEGs in which invasive and native genotypes showed higher basal-level expression in pairwise comparisons, respectively. Each cell shows the number and percentage of DEGs annotated with the GO term among all DEGs showing higher basal-level expression in the genotype specified by the column (red) compared to the genotype by the row (blue). Adjusted p-values of enrichment were calculated compared to a background of 41,595 reference gene models annotated with any GO term and represented as a color heatmap.

We aligned RNA-seq reads to the reference genome to explore the differences in basal transcriptome profiles between invasive and native genotypes and estimate the relative abundance of reference genes, as detailed in Methods (Supplementary Dataset 4). The three native genotypes showed overall 8.86% fewer reads aligned to the reference genome, compared to the three invasive genotypes, which showed an 88.92 ± 4.21 % alignment rate on average (Supplementary Table 5). When normalized to the total number of aligned reads, the distribution of estimated expression values did not show a noticeable bias towards either invasive or native genotypes (Supplementary Fig. 4). The leaf- and rhizome-derived transcriptomes were distinct from each other in a principal component analysis (PCA) (on PC1), while both showed additional separation between native and invasive genotypes (on PC2) (Fig 5c). In agreement with the PCA, the number of differentially expressed genes (DEGs) between invasive and native genotypes was much greater than DEGs detected between any other genotype comparisons (Fig. 5d). On average, 11.5 ± 1.0 % of the 64,857 reference gene models showed higher basal expression in the invasive genotypes compared to the native genotypes (Fig. 5d, upper right sections of the diagonal plots). Similarly, 12.1 ± 2.1% of gene models were more highly expressed in the native genotypes than in the invasive genotypes (Fig. 5d, lower left sections).

In the leaf samples, “response to stimulus” and “response to stress” were the largest representative GO terms showing significant enrichment among DEGs with higher basal expression in the invasive genotypes compared to the native genotypes in all cross-genotype comparisons (Supplementary Dataset 5). Interestingly, among child GO terms of these two GO terms, only those categorized as “response to biotic stimulus” and “defense response” showed significant enrichment in the invasive genotypes, but not “response to abiotic stress” (Supplementary Fig. 5). On average, 9.2% of all DEGs were annotated with “defense response” and showed higher basal expression in the invasive genotypes compared to 6.4% and 6.2% in the native genotypes and the background, respectively (Fig. 5e, upper panel). This bias towards higher a basal expression of genes associated with biotic stress and defense in the invasive genotypes was less clear in the rhizome tissue (Fig. 5e, lower panel). By contrast with the functional enrichment in defense responses observed for the invasive genotypes, the native genotypes were biased towards genes associated with “transmembrane transport” and its child GO terms, including “ammonium transport”, are represented by DEGs with higher basal expression in all cross-genotype comparisons (Fig. 5f and Supplementary Dataset 5). This transcriptomic signal was more prominent in leaf samples compared to rhizomes as similarly observed for the defense response.

### Divergence in invasive and native *P. australis* transcriptomic responses to biotic stress induced by fungal endophyte inoculation

We inoculated target genotypes with the fungal endophyte commonly isolated from *P. australis*, *Alternaria alternata* ^39^ (see Methods), to further investigate the transcriptomic signal established at basal expression contrasting the invasive genotypes from native genotypes biased towards biotic stress responses (Fig. 6, 1c, 5a). We compared 24 RNA-seq profiles generated for leaf and rhizome tissue harvested at pre-inoculation (basal expression) and post-inoculation (response to biotic stress) to deduce DEGs and their representative enriched functions in individual genotypes (see Methods, Supplementary Dataset 4). In general, more DEGs were significantly induced (Fig. 6a) than repressed (Fig. 6b) in response to the endophyte inoculation. The two invasive genotypes IOH1 and IMI1 showed more attenuated responses to endophyte inoculation compared to the three native genotypes. However, the invasive genotype IMI3 showed a response similar in magnitude to the native genotype NMI2 (Fig. 6a,b and Supplementary Dataset 6). Further, IMI3 and the three native genotypes shared a substantial number of endophyte-induced DEGs (Fig. 6a red boxes).

**Fig. 6.**
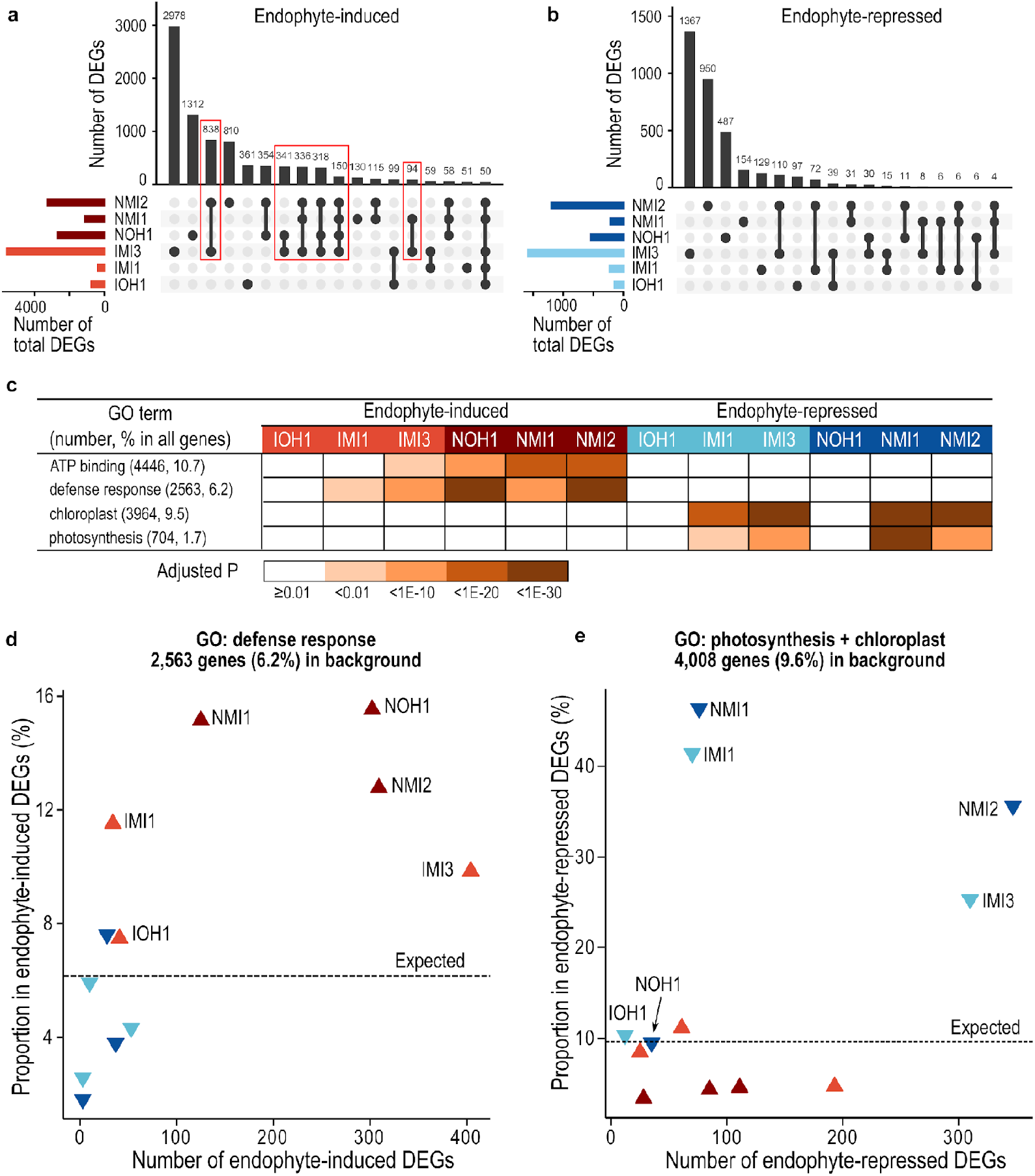
Transcriptome responses to *Alternaria alternata* fungal endophyte inoculation of invasive and native *P. australis* genotypes. **a-b,** Upset plots showing the number of shared and unique endophyte-induced (**a**) and repressed (**b**) DEGs among the six invasive and native genotypes. Red boxes show endophyte-induced DEGs shared between IMI3 genotype and a native genotype. **c**, The pattern of GO enrichment in six clones involving the largest number of endophyte-induced DEGs (represented by GO terms “ATP binding” and “defense response”) and endophyte-repressed DEGs (represented by GO terms “chloroplast” and “photosynthesis”). **d-e,** Number and percent proportion of endophyte-induced (upward triangles) or repressed (downward triangles) DEGs annotated with the GO term “defense response” (**d**) or either of the GO terms “photosynthesis” and “chloroplast” (**e**). The percent proportions in the entire *P. australis* gene models are marked with dashed lines as the expected value when there is no enrichment.

We searched for enriched functional associations between endophyte-induced and repressed genes in the invasive and native genotypes (Fig. 6c and Supplementary Dataset 6). In the native genotypes and IMI3 genotype, endophyte-induced genes were enriched in “ATP binding” and “defense response”, while endophyte-repressed genes were enriched in processes associated with photosynthesis (Fig. 6c). A closer inspection into these induced genes annotated under GO Molecular Function, “ATP-binding”, and GO Biological Process, “defense response”, revealed that many encode membrane receptor kinases known for roles in defense signaling (Supplementary Dataset 6).

The transcriptomic responses following endophyte inoculation may be reflective of local adaptations defined by each site selected for our study, compounding the broad distinctions we are able to deduce between native and invasive genotypes. For example, genotypes IMI3 and NMI2, while showing the highest number of induced DEGs annotated under defense response, also showed the highest number of repressed DEGs associated with photosynthesis compared to the rest of the genotypes, suggesting that the strong defense response concurrently down-regulated photosynthesis in these two genotypes (Fig. 6c-e). This potential trade-off related to photosynthesis seems to be less prominent in NOH1 where both the number and enrichment of photosynthesis-related endophyte-repressed DEGs were among the smallest compared to other genotypes. Despite genotypic variation in response to the endophyte inoculation, we found that in the native genotypes there was an overall stronger response in induced genes associated with biotic stress and defense, while these functions had higher basal expression in the invasive genotypes.

## Discussion

The genome of common reed, *Phragmites australis* reported here provides the first reference genome for the ecologically dominant and invasive model grass, spp. *australis* ^13,14^, and the first reference genome for the grass subfamily Arundinoideae (Fig. 1, 2, Table 1). Our genomic analyses, including non-reference allele frequency distributions and comparative genomics with other grasses, indicate that the reference genome was derived from a functionally diploid plant ^40^, despite *P. australis* being generally considered a tetraploid in North America ^25^. However, we found a previously unreported whole genome duplication (WGD) event, independent of three ancient WGD events known in the grasses ^36^, suggestive of *P. australis* being a paleo-tetraploid. The paleo-WGD event detected in the *P. australis* genome predates the divergence between spp. *australis* and *americanus*, but postdates the divergence from the Panicoideae (Fig. 3 and 4). As expected with substantial gene fractionation following WGD events, the *P. australis* genome lost up to 48% of its duplicated genes (Supplementary Fig. 2a), but retained over 14,005 duplicated genes. The retained duplicated genes were strongly enriched with transcription factors and other genes associated with regulatory processes. Genes encoding transcription factors were preferentially retained over other gene groups following paleo WGD events in pan-global species, such as *Arabidopsis thaliana ^41–43^*, and is thought to be a hallmark feature of WGD events underlying rapid diversification and global distribution of angiosperms^44,45^.

Adaptive innovations initiated at the genomic level further diversified at the transcriptome level can lead to distinct ecological fates. Our comparative analysis using native and invasive genotypes from the Great Lakes region indicates that gene expression associated with defense against biotic stress is primed in the invasive genotypes compared to native genotypes, and when a biotic stress response was induced using a fungal endophyte, the native genotypes were consistently more responsive (Fig. 5, 6). *P. australis* is known for its intraspecific ploidy polymorphism ^25,46^ coincident with its extreme success, being a globally recognized invasive species. Whether ploidy plays a deterministic role in facilitating invasiveness in *P. australis* remains debatable when categorical higher ploidy levels exclusively assigned to invasive genotypes are absent. However, as proposed by previous studies, it may predispose a species for invasive lifestyles adapted to a broad range of habitats potentially defined by both biotic and abiotic stresses ^47^. Therefore, exploring *P. australis* pangenomes in future studies should provide further insight into how ploidy polymorphism could add selective advantages to invasive genotypes of *P. australis* over their closely related non-invasive genotypes. Finally, our results provide a foundation for the development of novel species- and subspecies-specific genetic approaches for control of invasive *Phragmites* ^48^. There is widespread interest in controlling invasive *Phragmites* in many parts of its invasive range. Our genomic data help to identify particular genes that might be targeted in genetic control approaches such as RNAi, but our results also provide some caution in particular approaches if targeted genes are duplicated. While RNAi approaches have been widely explored for controlling invasive insect pests and plant pathogens ^49,50^, to our knowledge, no RNAi-based treatments have been developed to control problematic plants given the dearth of genomic data from invasive plant species. Future attempts at genetic control for invasive *Phragmites* should take into account that genetic variation exists in invasive *Phragmites* genotypes and that native genotypes can co-occur in the same habitats.

## Methods

### Plant materials

For genome sequencing, tillers and associated rhizome tissues were collected from a single *P. australis* clump of chloroplast haplotype M ^13,14^ at the Ottawa National Wildlife Refuge near Toledo, Ohio (Fig. 1c, marked with “G”) and propagated in a walk-in growth chamber as detailed in Supplementary Methods. For transcriptome analyses, we collected three additional invasive and three native genotypes from four sites around the Great Lakes in Michigan and Ohio, U.S. (Fig. 1c and Fig. 5a). Native genotypes were readily distinguished from invasive genotypes by their smaller stature and thinner tillers with distinctive reddish coloration at the nodes (Fig. 1e, f, g). Both native and invasive genotypes were confirmed by chloroplast haplotype sequences as previously described ^13,51^. *Phragmites* plants were grown in the growth chamber and subjected to endophyte inoculation treatments before RNA isolation for RNA-seq analyses, as detailed in Supplementary Methods.

### Genome sequencing, assembly, and annotation

Genomic DNA was isolated from leaf tissue using a 2% CTAB extraction protocol ^52^, converted to SMRTbell^TM^ libraries (Pacific Bioscience, Menlo Park, CA) with 20-Kb target insert size, and sequenced for continuous long reads (CLR) using a PacBio RSII single-molecule real-time sequencing platform. After post-processing (SMRT Analysis v2.3, Pacific Bioscience), total 4.79 million reads (mean length 8.80 Kbp, N50 13.11 Kbp, and total 42.19 Gbp) were assembled into contigs using Canu assembler (v. 1.4) ^31^, with the target genome size set to 1 Gbp, generating the reference genome assembly. In addition, the same genomic DNA samples were converted to TruSeq DNA paired-ends libraries (Illumina, San Diego, CA) for whole-genome shotgun sequencing, analyzed for 300 cycles on a NextSeq platform (Illumina), and used to estimate the genome ploidy levels (Supplementary Methods and Supplementary Fig. 1).

For genome annotation, we first used RepeatModeler v. 1.0.8 and RepeatMasker v. 4.0.9 (http://www.repeatmasker.org/) to detect transposable elements and repetitive sequences in the assembled *P. australis* contigs. *De novo*-detected repetitive sequences were combined with known monocot repeat sequences in RepBase (v. 23.07; http://www.girinst.org/) to mask repeats in *P. australis* contigs. Protein-coding gene models on the repeat-masked genome were predicted using the MAKER (v. 2.31.10) ^53^ with *P. australis* BUSCO-trained parameters for *ab initio* gene model prediction^32^ and hints from *P. australis* transcriptomes *de novo* assembled using Trinity (v. 2.1.1) ^54^ as well as from homologs from *Sorghum* (Phytozome ID:454) and *Brachypodium* (Phytozome ID:314). In addition, we performed a reference-guided transcriptome assembly using StringTie (v. 2.0.1) ^55^ to report putative isoform models. The gene model encoding the longest open reading frame (ORF) was selected as the representative for each of the 64,857 protein-coding gene loci.

### Comparative and functional analyses

We used BUSCO (v. 3) ^32^ to assess the completeness of the representative *P. australis* protein-coding gene models in comparison with grass genomes available in Monocots PLAZA (v. 4.5) ^33^ (Supplementary Table 2). Maximum-likelihood species trees were estimated by OrthoFinder (v. 2.2.7) ^56^ and RAxML (v. 8.2) ^57^ using concatenated alignment of 782 sequences. We selected five monocot genomes for in-depth comparative analyses, based on their availability of recently updated gene models with >90% BUSCO scores and lack of genome duplication more recent than the ρ event. MCscan (as implemented in JCVI v. 1.1.7) ^58^ was used to compare syntenic depths between *P. australis* and other monocot genomes. SynMap ^59^ and CLfinder pipeline ^60^ detected co-linear paralog and ortholog pairs within the *P. australis* genome and between *P. australis* and monocot genomes, respectively. We estimated synonymous substitution rates at 4-fold degenerate sites using codeml ^61^ as described previously ^60^. We detected ortholog groups among representative gene models of *P. australis* and other monocot genomes using OrthoFinder (v. 2.2.7) ^56^ and MMseqs2 ^62^, and subsequently identified *P. australis* ortholog groups that are “Conserved” and “Duplicated” in ortholog copy numbers compared to other monocot species, as detailed in Supplementary Methods.

Gene ontology (GO) annotation was transferred to the *Phragmites* representative gene models based on sequence similarities to plant proteins with GO annotations as of January 1^st^, 2020 in the GO consortium (http://geneontology.org/). In short, *Phragmites* protein sequences were compared with reference sequences with a GO annotation using MMseqs2 with the maximum sensitivity (−s 7.5). If the protein alignment covers minimum 30% of both the query and subject sequences, the GO annotation was transferred to the *Phragmites* protein. We used BiNGO ^63^ to detect GO terms enriched in *Phragmites* gene models. GO terms were further clustered and summarized using GOMCL ^37^.

### Endophyte treatment, transcriptome assembly, and RNA-seq analyses

Invasive and native genotypes of *P. australis* were propagated from rhizome cuttings, grown in a growth chamber for 60 days, and subjected to *Alternaria alternata* (accession KT923239) ^39^ fungal endophyte inoculation and RNA-Seq, as detailed in Supplementary Methods. RNA-seq reads were filtered of adapter sequences using FASTP ^64^. Filtered reads were aligned to the reference genome using HISAT2 (v. 2.2) and expression of each gene model was estimated with StringTie (v. 2.0.1) with default parameters ^65^. Differentially Expressed Genes (DEGs) were identified based on a minimum 2-fold difference in expression levels, with adjusted p-values <0.05 (estimated by DESeq2 ^66^), between pairs of genotypes or between endophyte-inoculated and control samples within each genotype. In addition, putative protein-coding transcript sequences were obtained from filtered RNA-Seq reads by the Trinity (v. 2.1.1) and TransDecoder (v. 5.5.0) pipeline ^54^ (Supplementary Table 4) and used for estimating the phylogenetic relationship of *P. australis* subspecies and genotypes using the Agalma pipeline (v. 2)^67^ (Fig. 3d and 5b).

## Supporting information

All Supplemental Materials

## Data availability

All raw and assembled sequence data are deposited to NCBI under BioProject PRJNA705976. In addition, the reference genome sequence, gene models, and representative RNA-Seq tracks are available in the CoGe database with the Genome ID 59768, for browsing and other CoGe-embedded comparative analyses ^68^.

## Acknowledgements

The authors would like to thank The U.S. Fish and Wildlife Service and Michigan Department of Environment, Great Lakes and Energy for access to study sites and the Great Lakes Restoration Initiative for financial support. We thank Doug Rusch and Ram Podicheti and the staff of the Indiana University Center for Genomics and Bioinformatics for their help in initial genome and transcriptome analyses, Matt Filapek for lab assistance, and the Indiana University Greenhouse staff for their assistance with growing and propagating *Phragmites*. We thank Dr. Wesley Bickford for data collection and analysis support and Drs. Ping Gong, Kathleen Ferris, and Simon Barak for their thoughtful comments. This research was supported by the United States Geological Survey Cooperative Agreement G18AC00373 to KC. MD and DHO acknowledge the support of National Science Foundation awards MCB-1616827, NSF-IOS-EDGE-1923589, and the Next-Generation BioGreen21 Program of Republic of Korea (PJ01317301). CW was supported by an Economic Development Assistantship award from Louisiana State University. The authors also acknowledge the LSU High Performance Computing services for providing computational resources needed for data analyses.

## Author contributions

K.C., M.D., and K.K. conceived and designed the experiments and obtained the funding. P.T., K.C., and K.K. conducted data collections and related experiments. D-H.O., C.W, and MD designed the bioinformatics work flows and performed computational analyses. D-H.O and Q.Q. generated figures. All authors contributed to drafting and finalizing the manuscript.

## Contents of Supplementary Materials (provided separately)

### Supplementary Methods

**Supplementary Table 1** | Proportion of transposable elements in the *Phragmites australis* reference genome

**Supplementary Table 2** | Monocot genomes for comparative analyses

**Supplementary Table 3** | Comparison of orthologous gene copy numbers among *P. australis* and five monocot genomes

**Supplementary Table 4** | *De novo* assembled *P. australis* transcriptomes

**Supplementary Table 5** | Alignments of RNA-seq reads derived from invasive and native genotypes to the *P. australis* reference genome

**Supplementary Fig. 1** | Distribution of non-reference allele frequencies indicates a functionally diploid genome of the *P. australis* genotype used for the reference genome assembly.

**Supplementary Fig. 2** | Properties of synteny blocks identified within the *P. australis* genome.

**Supplementary Fig. 3** | The fifty largest synteny blocks detected within the *P. australis* genome.

**Supplementary Fig. 4** | RNA-seq analysis comparing basal-level expression among native and invasive *P. australis* genotypes.

**Supplementary Fig. 5** | An example network of Gene Ontology (GO) terms enriched among *P. australis* reference genes with higher basal-level expression in an invasive genotype.

The following datasets are available at the USGS repository (https://doi.org/10.5066/P91XSF3):

**Supplementary Dataset 1.** *P. australis* version 1.0 reference gene models and syntelog pairs

**Supplementary Dataset 2.** Ortholog groups identified among *P. australis* and five monocots

**Supplementary Dataset 3.** GO clusters enriched among *P. australis* ortholog groups showing conserved or increased gene copy numbers due to *P. australis*-specific duplications

**Supplementary Dataset 4.** RNA-Seq raw read counts and expression rank percentiles for *P. australis* gene models for invasive and native genotypes.

**Supplementary Dataset 5.** RNA-Seq results comparing basal-level expression of *P. australis* genes among invasive and native genotypes

**Supplementary Dataset 6.** GO terms enriched among *P. australis* genes with significantly different basal-level expression among invasive and native genotypes.

**Supplementary Dataset 7.** RNA-Seq results for endophyte-responses of *P. australis* genotypes

**Supplementary Dataset 8.** GO terms enriched among *P. australis* genes whose expression were significantly induced or repressed by endophyte inoculation.

