## Supplementary material for "Novel genome characteristics contribute to the invasiveness of *Phragmites australis* (common reed)": All Supplemental Materials

**Supplementary Materials for: Novel genome characteristics contribute to the invasiveness of *Phragmites australis* (common reed)**

Dong-Ha Oh, Kurt P. Kowalski, Quynh N. Quach, Chathura Wijesinghege, Philippa Tanford, Maheshi Dassanayake, and Keith Clay

### Supplementary Methods

Propagation of *Phragmites australis* genotypes in growth chambers

Phragmites invasive and native genotypes were propagated from rhizome cuttings at Indiana University and maintained in large clay pots in the Indiana University Greenhouse. Rhizome cuttings with developing buds (3-10cm in length) were planted in potting soil in round plastic pots (9cm height, 11cm diameter) and watered as needed. Potted cuttings were kept in the greenhouse until transfer to a Percival Scientific Model WE-1012 Environmental Controller walk-in growth chamber kept at 70°F and 70% relative humidity setting with a 14:10 hour light:dark cycle for approximately 60 days before further manipulations.

Endophyte treatment and RNA sample preparation

**Endophyte treatments**. Replicate plants from all genotypes were inoculated with an *Alternaria alternata* fungal endophyte (Accession KT923239), one of the most abundant fungal OTUs isolated from *Phragmites* [39], to potentially elicit *Phragmites* transcriptional responses. Fungal inoculum from stock cultures was used to inoculate four flasks with 50mL liquid culture medium (M102) [70] in 1L distilled $H_2O$, which were then cultured for 4 days on a Braun Melsungen AG auto shakeR Type 886013 shaker table at 120 rpm. One tiller per plant was inoculated using three methods. First, a sterile 1ML needle-tipped syringe covered in *Alternaria alternata* mycelium was inserted into the tiller above the first node. Second, a sterile wooden toothpick overgrown with mycelium from liquid culture was inserted above the second node and left in place overnight. Finally, three sequential leaves on the inoculated tiller were rubbed with sterile chromatography sand to create small wounds to the leaf epidermis, and the entire plant was treated with inoculum spray. Spray inoculum was prepared by decanting the contents of four 50mL liquid culture flasks into 50 mL centrifuge tubes (~30mL/tube). Sterile DI water was added to each tube and the contents were thoroughly mixed by shaking, centrifuged, and the supernatant discarded. This process was repeated a second time and 500mL DI water was added to the rinsed mycelium and blended for 30 seconds in a sterile blender, and transferred to a 550mL plastic spray bottle for several hours before application. Abundant mycelium was visible on a microscope slide at 200X magnification following a single spray, but few or no fungal spores were present. Control plants received identical treatments but with sterile needles, sterile toothpicks, and a 2% liquid culture media solution in sterile DI water. Treated plants were then covered and secured in a plastic bag overnight to maintain high humidity. The growth chamber was kept dark for 36 hours after inoculation before resuming a 14:10 hour light/dark cycle.

**Plant tissue sampling and inoculum isolation**. Two inoculated and two control plants per genotype were sampled eight days after inoculation treatment applications for transcriptome analysis. Soil was washed away in tap water and the plant was blotted dry before the treated tillers and attached rhizome were processed. We sampled the first 2cm of rhizome directly connected to the infected tiller. If more rhizome tissue was present, 0.5cm rhizome sections were taken every 2cm or until a total of 5cm was sampled (2cm + six 0.5cm samples). The rhizome sections were cut in half lengthwise with a clean razor blade to replicate samples for RNA extraction. Three leaves from the infected tiller were each cut into quarters after a sample was taken to confirm inoculation. The top and bottom quarter of each leaf were pooled to replicate samples for RNA extraction.

Leaf tissue samples were taken from the three inoculated leaves per plant rubbed with sand to confirm the effectiveness of inoculation. A $1cm^2$ leaf section from each leaf was cut into small fragments, placed in small plastic tissue cassettes and rinsed for 20 minutes under running tap water individually. Leaf fragments within the tissue cassettes were then washed for 1 minute in 95% ethanol, 3 minutes in 0.18% sodium hypochlorite, 1 minute in 95% ethanol, and 2 minutes in sterile DI water following Christian et al.[71]. Each leaf fragment was removed from the tissue cassette, air-dried on a sterile paper towel and placed on the surface of a corn meal agar (CMA) petri plate supplemented with 25mg tetracycline/L for 5 minutes to confirm the efficacy of surface sterilization. Each fragment was then cut in half with a sterile razor to expose the plant interior and plated on another CMA + tetracycline plate to determine if subsequent fungal growth corresponds to the original *Alternaria alternata* inoculum.

**RNA Extraction**. Tissue samples were grounded in liquid nitrogen in porcelain mortars and pestles while the mortar was resting in dry ice. Mortars and pestles were previously wrapped in aluminum foil and baked in a drying oven at 180°C for six hours to destroy RNase and then pre-chilled in a -20°C freezer. Grounded tissue and liquid nitrogen was decanted into a 2mL microcentrifuge tube and liquid nitrogen was allowed to evaporate. ~1mL of RLT buffer with 10µl 2-mercaptoethanol per mL was then added. Each sample was promptly vortexed and put on ice until transfer to long-term storage at -80°C. Between samples (except replicates), mortars and pestles were washed with soap and water, sprayed with RNase AWAY (Molecular BioProducts, Inc.), and re-chilled. Most samples were processed within ten minutes of cutting.

RNA was extracted using the RNeasy Plant Mini Kit (Qiagen) following the manufacturer's protocol with the following minor modifications. Samples were thawed on ice, and centrifuged for 15-30 seconds at 10,000xg to pellet plant tissue and allow for pipetting of the supernatant. Then, 300 µl of sample was added to the Qiashredder column (up to ~800ul for samples with lower concentration). Extracts were stored at -80°C in RNase-free tubes.

**RNA Library preparation and sequencing.** cDNA libraries were prepared with the Illumina Stranded mRNA Prep Kit (Illumina) following the standard protocol with half volume reactions and 12 PCR amplification cycles in the library enrichment step. Libraries from samples of low concentration (<500ng material available) were constructed from 25ul of input material regardless of concentration. Final library quality and concentration for pooling was assessed with Agilent D1000 ScreenTape (Agilent). Pooled libraries were sequenced through a NextSeq 75 cycle high-output run with paired-end read on an Illumina NextSeq500 at the Center for Genomics and Bioinformatics at Indiana University.

<u>Determination of non-reference allele frequencies in single nucleotide polymorphism (SNP) loci.</u>

Before gene model prediction, we aligned paired-end short reads derived from the same genomic DNA samples used for the reference genome assembly, to the reference genome sequence using bowtie2 (v. 2.2.6) [72]. We then determined non-reference allele frequencies in genomic regions with homologies to Benchmarking Universal Single-Copy Ortholog (BUSCO) sequences [73] using mpileup command of samtools (v. 1.8) with a Phred quality score cutoff of 20 [74] (Supplementary Fig. 1). To determine the genomic regions harboring putative BUSCOs, we first detected transcripts homologous to the 956 BUSCO HMM profiles [73] from the "Invasive combined" transcriptome (Supplementary Table 4). These transcripts were mapped to the genomic regions using GMAP (v. 2012-04-21) [75].

Identification of "Conserved" and "Duplicate" groups among *P. australis* reference gene models.

Using OrthoFinder (v. 2.2.7) [56] and MMseqs2 [62], we identified ortholog groups (OGs) among deduced protein sequences from the primary reference gene models of *P. australis* and five comparator monocot species. For each OG, we calculated the ratio of ortholog copy number ($R_{CN}$) for each species, as defined as the ortholog copy number in a species divided by the mean ortholog copy number of all other species (Fig. 4a). OGs with the standard deviation of ortholog copy number across all species exceeding 10% of the total number of orthologs were excluded since such OGs tend to include lineage-specific expansion of transposable element-like genes whose copy numbers are less relevant with the *P. australis*-specific whole genome duplication (WGD) event (Fig. 3 and Supplementary Dataset 2). Among *P. australis* OGs, we identified "Conserved" and "Duplicated" groups whose $R_{CN}$ values are between 0.75 to 1.25 ("Conserved") and larger than 1.5 ("Duplicated"), respectively (Fig. 4a and Supplementary table 2), representing *P. australis* orthologs whose copy numbers have been fractionated to the level similar to other monocot species ("Conserved") or retained as duplicated after the WGD event.

**References** (numbered in continuation from the main text)


70. Bacon, C. W. *Biotechnology of Endophytic Fungi of Grasses*. (CRC Press, 2018).
71. Christian, N., Herre, E. A. & Clay, K. Foliar endophytic fungi alter patterns of nitrogen uptake and distribution in *Theobroma cacao*. *New Phytol.* **222**, 1573–1583 (2019).
72. Langmead, B. & Salzberg, S. L. Fast gapped-read alignment with Bowtie 2. *Nat. Methods* **9**, 357–359 (2012).
73. Chan, K.-L. *et al.* Seqping: gene prediction pipeline for plant genomes using self-training gene models and transcriptomic data. *BMC Bioinformatics* **18**, 1426 (2017).
74. Li, H. A statistical framework for SNP calling, mutation discovery, association mapping and population genetical parameter estimation from sequencing data. *Bioinformatics* **27**, 2987–2993 (2011).
75. Wu, T. D. & Watanabe, C. K. GMAP: a genomic mapping and alignment program for mRNA and EST sequences. *Bioinformatics* **21**, 1859–1875 (2005).

**Supplementary Table 1 | Proportion of transposable elements in the *Phragmites australis* reference genome**

| Category | Number of elements | Length occupied (nt) | Percent occupied (%) |
|---|---|---|---|
| SINEs | 8,937 | 1,643,956 | 0.14 |
| LINEs | 33,384 | 19,841,853 | 1.74 |
| LTR elements | 326,358 | 415,123,549 | 36.42 |
| DNA elements | 299,783 | 130,261,850 | 11.43 |
| Unclassified | 214,476 | 73,623,178 | 6.46 |
| Total interspersed repeats | | 640,494,386 | 56.19 |
| Total sequence except gaps | | 1,139,927,050 | 100.00 |

**Supplementary Table 2 | Monocot genomes for comparative analyses**

| Species | Version | Source | BUSCO (%) [b] | Protein-coding gene loci |
|---|---|---|---|---|
| *Ananas comosus* [a] | 3 | Phytozome (ID:321) | C:91.8[S:87.4,D:4.4],F:3.6,M:4.6 | 27,024 |
| *Brachypodium distachyon* [a] | 3.1 | Phytozome (ID:314) | C:99.5[S:98.3,D:1.2],F:0.5,M:0.0 | 34,310 |
| *Cenchrus americanus* | 1.0 | PLAZA (4.5) | C:82.3[S:79.8,D:2.5],F:11.3,M:6.4 | 38,579 |
| *Hordeum vulgare* | 2.44 | IBSC | C:84.8[S:80.1,D:4.7],F:3.9,M:11.3 | 36,760 |
| *Leersia perrieri* | N/A | Ensembl (r42) | C:92.5[S:90.8,D:1.7],F:3.7,M:3.8 | 29,176 |
| *Oryza sativa* Japonica [a] | 7 | Phytozome (ID:323) | C:95.1[S:93.7,D:1.4],F:3.4,M:1.5 | 42,189 |
| *Oropetium thomaeum* | 1.0 | Phytozome (ID:386) | C:81.2[S:78.4,D:2.8],F:13.7,M:5.1 | 28,437 |
| *Phragmites australis* [a] | 1.0 | This study | C:93.3[S:56.6,D:36.7],F:4.9,M:1.8 | 64,857 |
| *Phyllostachys edulis* | 1.0 | NCGR | C:73.5[S:61.1,D:12.4],F:13.7,M:12.8 | 31,987 |
| *Sorghum bicolor* [a] | 3.1.1 | Phytozome (ID:454) | C:99.3[S:97.8,D:1.5],F:0.4,M:0.3 | 34,129 |
| *Setaria italica* [a] | 2.2 | Phytozome (ID:312) | C:98.4[S:96.7,D:1.7],F:1.1,M:0.5 | 34,584 |
| *Saccharum spontaneum* | v20190103 | PLAZA (4.5) | C:92.4[S:22.7,D:69.7],F:3.4,M:4.2 | 83,826 |
| *Triticum aestivum* | 1.1.44 | IWGSC | C:99.6[S:1.1,D:98.5],F:0.1,M:0.3 | 107,327 |
| *Zea mays* B73 | 4 | AGP | C:94.3[S:84.9,D:9.4],F:3.9,M:1.8 | 39,421 |
| *Zoysia japonica nagirizaki* | 1.1 | PLAZA (4.5) | C:45.9[S:41.0,D:4.9],F:28.9,M:25.2 | 59,271 |

[a] Selected for comparative analyses in Fig. 3 and Fig. 4.

[b] Based on "embryophyta odb10" with total 1,375 Benchmarking Universal Single-Copy Orthologs (BUSCOs), shown are percentages of complete (C) single-copy (S), duplicated (D), fragmented (F), and missing (M) BUSCOs.

**Supplementary Table 3 | Comparison of orthologous gene copy numbers among *P. australis* and five monocot genomes**

| Copy number ratio compared to others ($R_{CN}$)[a] | Number of Ortholog groups (OGs) | | | | | |
|---|---|---|---|---|---|---|
| | *A.comosus* | *B. distachyon* | *O. sativa* | *P. australis* | *S. bicolor* | *S. italica* |
| $0.0 \leq R_{CN} \leq 0.125$ | 1136 | 59 | 73 | 92 | 20 | 26 |
| $0.125 < R_{CN} \leq 0.25$ | 44 | 3 | 2 | 0 | 1 | 1 |
| $0.25 < R_{CN} \leq 0.375$ | 145 | 23 | 13 | 11 | 12 | 5 |
| $0.375 < R_{CN} \leq 0.5$ | 892 | 248 | 194 | 103 | 133 | 80 |
| $0.5 < R_{CN} \leq 0.625$ | 798 | 751 | 696 | 301 | 673 | 611 |
| $0.625 < R_{CN} \leq 0.75$ | 573 | 823 | 800 | 259 | 809 | 741 |
| $0.75 < R_{CN} \leq 0.875$ | 2600 | 3061 | 3052 | 851 | 3039 | 2985 |
| $0.875 < R_{CN} \leq 1.0$ | 3134 | 3539 | 3568 | 3051 | 3613 | 3603 |
| $1.0 < R_{CN} \leq 1.125$ | 139 | 395 | 433 | 278 | 461 | 495 |
| $1.125 < R_{CN} \leq 1.25$ | 414 | 1562 | 1558 | 1463 | 1661 | 1680 |
| $1.25 < R_{CN} \leq 1.375$ | 55 | 45 | 64 | 96 | 56 | 78 |
| $1.375 < R_{CN} \leq 1.5$ | 284 | 272 | 295 | 475 | 290 | 325 |
| $1.5 < R_{CN} \leq 1.625$ | 14 | 14 | 18 | 45 | 26 | 15 |
| $1.625 < R_{CN} \leq 1.75$ | 430 | 181 | 205 | 943 | 184 | 260 |
| $1.75 < R_{CN} \leq 1.875$ | 29 | 29 | 28 | 226 | 30 | 41 |
| $1.875 < R_{CN} \leq 2.0$ | 371 | 109 | 106 | 2118 | 103 | 149 |
| $2.0 < R_{CN} \leq 2.125$ | 2 | 3 | 2 | 18 | 2 | 5 |
| $2.125 < R_{CN} \leq 2.25$ | 21 | 13 | 21 | 246 | 16 | 25 |
| $2.25 < R_{CN} \leq 2.375$ | 0 | 4 | 2 | 33 | 4 | 5 |
| $2.375 < R_{CN} \leq 2.5$ | 66 | 14 | 18 | 440 | 14 | 18 |
| $2.5 < R_{CN} \leq 2.625$ | 0 | 0 | 0 | 0 | 0 | 0 |
| $2.625 < R_{CN} \leq 2.75$ | 0 | 0 | 0 | 8 | 0 | 0 |
| $2.75 < R_{CN} \leq 2.875$ | 1 | 0 | 0 | 91 | 1 | 0 |
| $2.875 < R_{CN}$ | 0 | 0 | 0 | 0 | 0 | 0 |

[a] Orthologous gene copy number ($R_{CN}$) was calculated for each species in an OG, by dividing the ortholog in the species by the mean copy number in all other species. The number of OGs in different $R_{CN}$ range is shown for all species. For *P. australis*, OGs in "Conserved" and "Duplicated" groups are shaded in orange and skyblue, respectively. See Supplementary Methods for details.

**Supplementary Table 4 | *De novo* assembled *P. australis* transcriptomes**

| Transcriptome | # ORFs | min (nt) | median (nt) | mean (nt) | N50 (nt) | max (nt) | sum (nt) |
|---|---|---|---|---|---|---|---|
| Invasive combined [a] | 97,076 | 123 | 678 | 895 | 1,230 | 14,250 | 86,956,037 |
| Native leaf[b] | 49,224 | 255 | 672 | 919 | 1,155 | 12,597 | 45,275,301 |
| Native rhizome | 56,916 | 258 | 768 | 998 | 1,263 | 13,053 | 56,843,553 |
| IOH1 leaf[c] | 34,131 | 255 | 699 | 950 | 1,191 | 12,600 | 32,448,375 |
| IOH1 rhizome | 27,866 | 258 | 744 | 1,009 | 1,290 | 8,745 | 28,125,045 |
| IMI1 leaf | 35,114 | 255 | 702 | 956 | 1,209 | 12,600 | 33,576,678 |
| IMI1 rhizome | 44,499 | 258 | 804 | 1,040 | 1,308 | 13,548 | 46,311,834 |
| IMI3 leaf | 34,890 | 258 | 717 | 964 | 1,212 | 13,053 | 33,665,640 |
| IMI3 rhizome | 39,165 | 261 | 810 | 1,031 | 1,281 | 12,648 | 40,404,453 |
| NOH1 leaf | 39,466 | 255 | 690 | 939 | 1,185 | 12,918 | 37,085,379 |
| NOH1 rhizome | 46,233 | 255 | 798 | 1,024 | 1,275 | 12,762 | 47,376,243 |
| NMI1 leaf | 42,354 | 261 | 708 | 963 | 1,218 | 12,795 | 40,798,848 |
| NMI1 rhizome | 48,766 | 261 | 789 | 1,030 | 1,305 | 13,053 | 50,252,232 |
| NMI2 leaf | 43,633 | 258 | 690 | 939 | 1,185 | 12,777 | 40,976,049 |
| NMI2 rhizome | 37,619 | 255 | 735 | 994 | 1,266 | 15,036 | 37,424,133 |

Stats for open reading frames (ORFs) are shown for all *de novo* assembled transcriptomes. [a] RNA-seq reads (150nt x2) from leaf, shoot, and rhizomes of the same clone used for genome assembly were assembled with Trinity (v. 2.4.0) and SPAdes (v. 3.10.0) and consolidated using the evigene pipeline. This transcriptome was used as the "EST hints" for reference gene model annotation. [b] Half of all RNA-seq reads (37nt x2) from the three native genotypes were combined and assembled as described in Methods. The Native rhizome transcriptome was used as the representative of *P. australis* native subspecies (ssp. *americanus*) in Fig. 3c and 3d. Using the Native leaf transcriptome produced similar results. [b] All RNA-seq reads (37nt x2) from each tissue were used to assemble transcriptomes for each genotype, as described in Methods. The rhizome transcriptomes represented the six genotypes in Fig 5b. The leaf transcriptomes produced a near identical tree.

**Supplementary Table 5 | Alignments of RNA-seq reads derived from invasive and native genotypes to the *P. australis* reference genome**

| ID | Genotype | Total million read pairs (mean ± s.d.) [a] | % overall alignment rate (mean ± s.d.) [a] |
|---|---|---|---|
| IOH1 | Invasive | 29.55 ± 4.34 | 89.82 ± 3.44 |
| IMI1 | Invasive | 28.48 ± 3.26 | 89.52 ± 3.02 |
| IMI3 | Invasive | 28.88 ± 2.82 | 87.43 ± 5.36 |
| NOH | Native | 29.05 ± 3.92 | 78.32 ± 8.72 |
| NMI1 | Native | 28.36 ± 3.17 | 80.65 ± 1.21 |
| NMI2 | Native | 30.84 ± 6.56 | 81.21 ± 1.26 |

[a] Mean and standard deviation (s.d.) for total 32 RNA-seq samples per genotype, i.e. two tissues (leaf and rhizome) x two treatment (control and inoculated with endophytes) x eight replicates

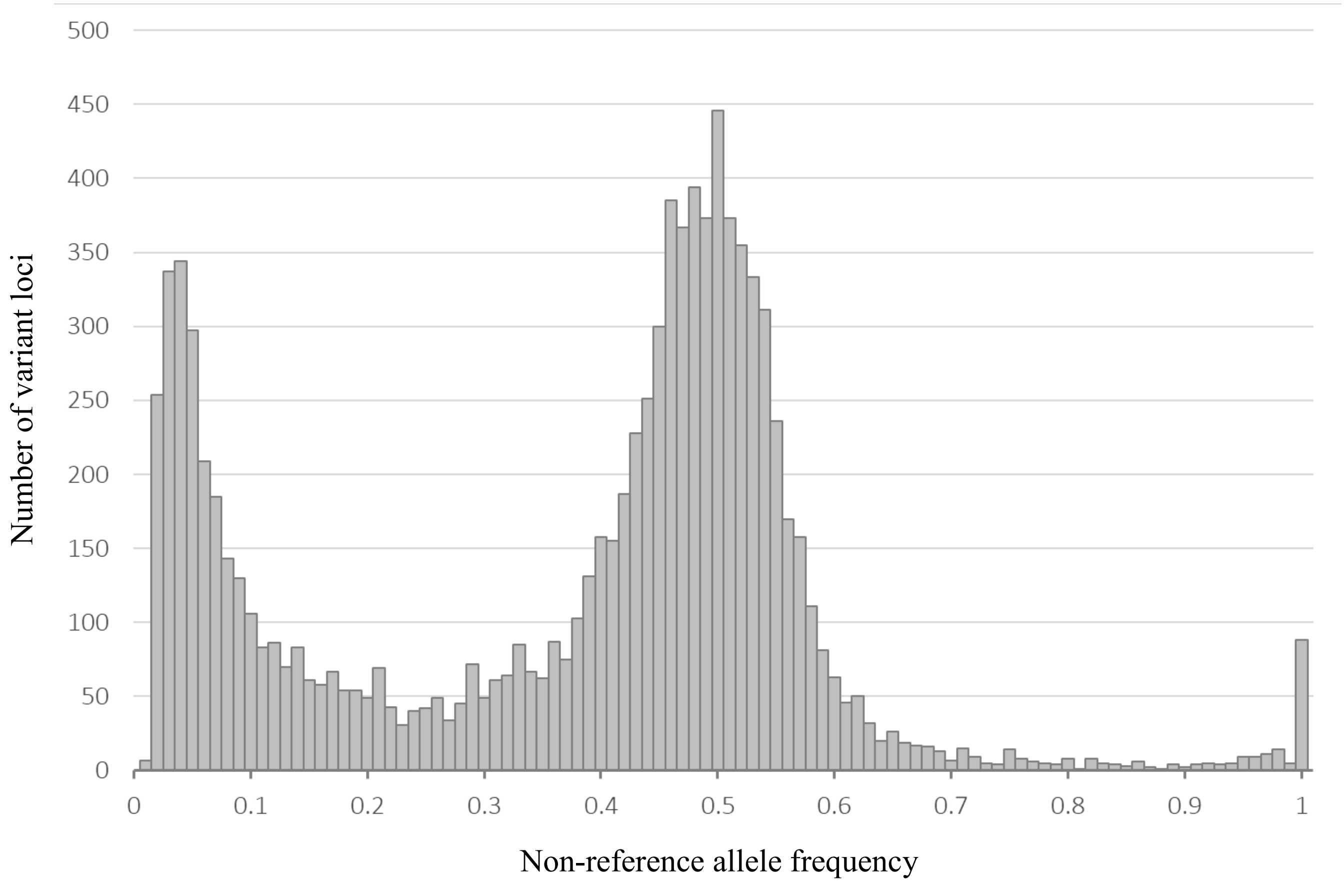


**Supplementary Fig. 1 | Distribution of non-reference allele frequencies indicates a functionally diploid genome of the *P. australis* genotype used for the reference genome assembly.** We identified total 9,835 single nucleotide polymorphisms in total 3,780,373-bp genomic regions harboring similarities with 699 BUSCOs and plotted the distribution of non-reference allele frequencies (see Supplementary Methods for details). The single major peak at allele frequency 0.5 coincided with the expectation from a diploid genome.

**a**

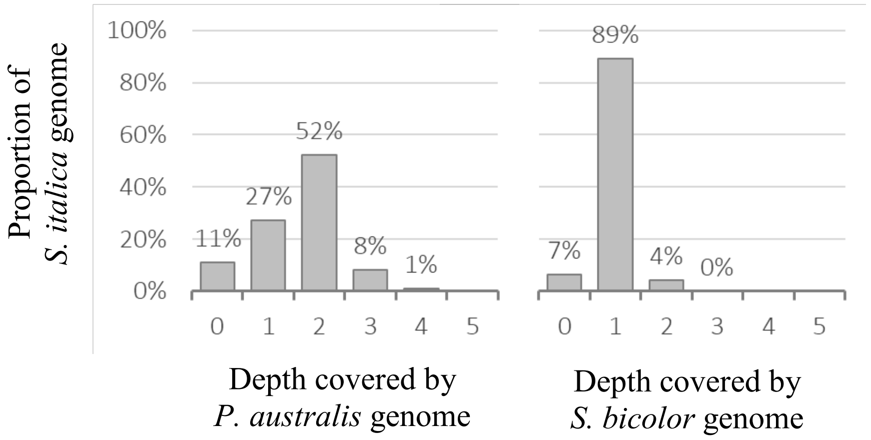


**b**

| Contig size (# protein-coding gene loci) | Number of contigs | Number (%) of contigs with a synteny block |
|---|---|---|
| ≥5 | 3551 | 1357 (38.2) |
| ≥10 | 1549 | 1035 (66.8) |
| ≥20 | 590 | 541 (91.7) |
| ≥50 | 107 | 106 (99.1) |
| ≥100 | 22 | 22 (100) |

**c**

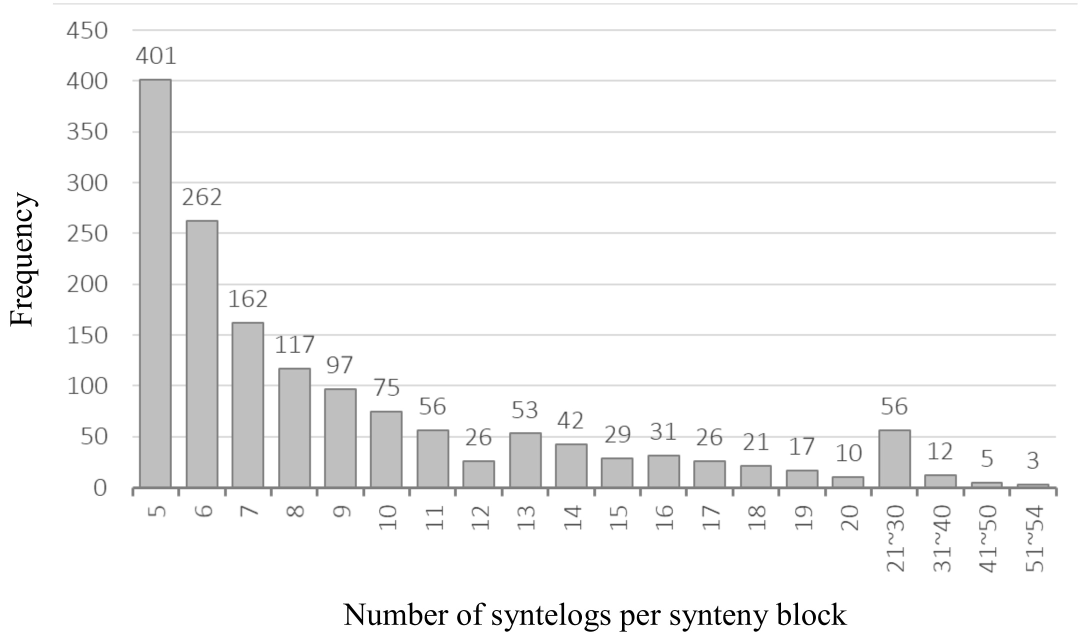


**Supplementary Fig. 2 | Properties of synteny blocks identified in the *P. australis* genome. a,** Comparison of the *Setaria italica* genome proportions covered by different depths with *Phragmites australis* and *Sorguhm bicolor* genomes, respectively. Total 52% of *S. italica* genes were paired with two *P. australis* co-linear orthologs as exemplified in Fig. 3b, while most

(>89%) of *S. italica* genes were 1:1 matched with a *S. bicolor* co-linear ortholog. **b,** Number and proportion of *P. australis* draft genome contigs that include a synteny block with minimum five co-linear paralogs. **c,** Size distribution of synteny blocks. Synteny blocks, detected using SynMap, are defined as a stretch of minimum five co-linear paralogs (i.e. syntelog) in other parts of the same genome. See the CoGe SynMap description (https://genomevolution.org/wiki/index.php/SynMap) for details.

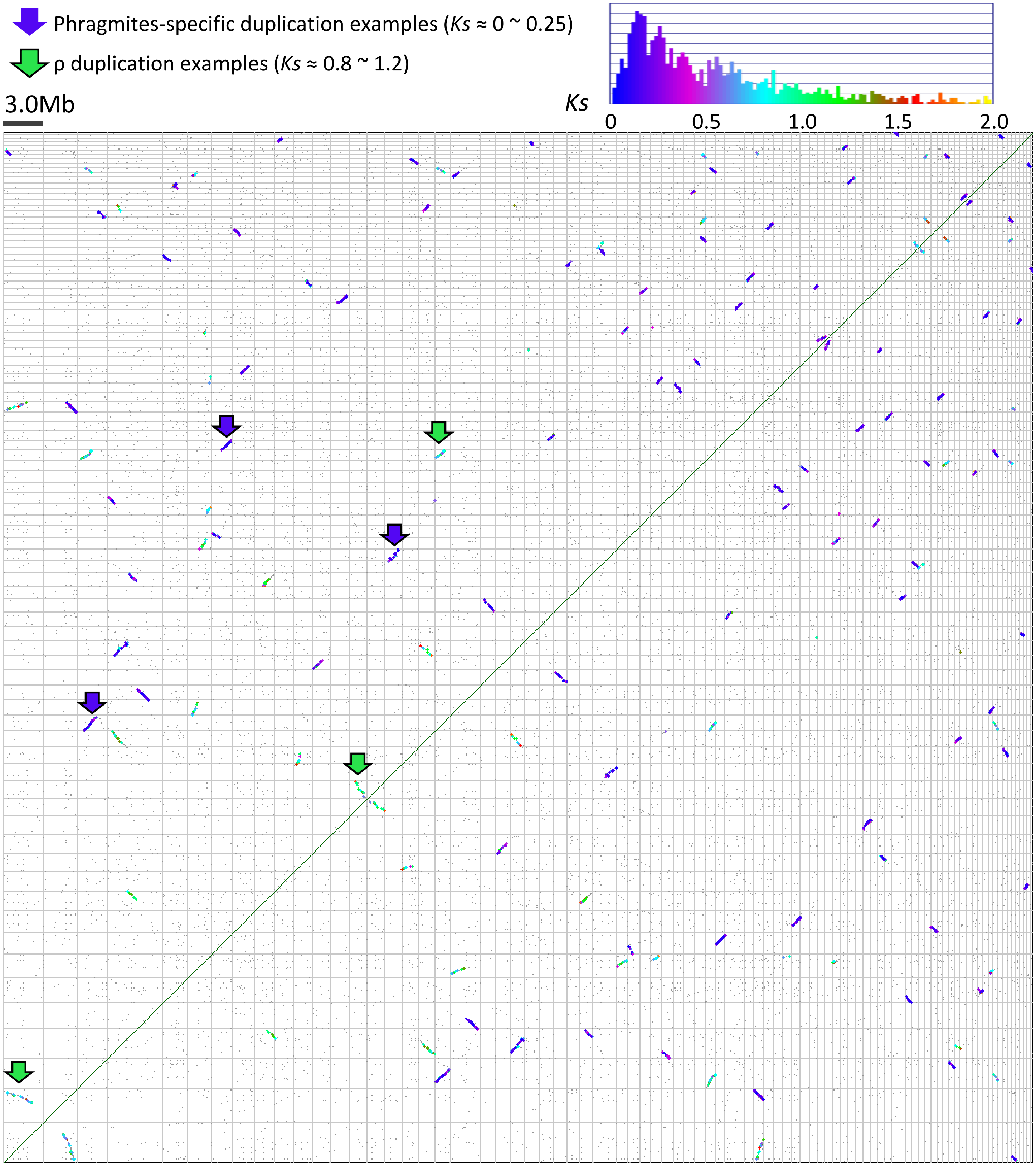


**Supplementary Fig. 3 | The fifty largest synteny blocks detected within the *P. australis* genome.** *P. australis* genome contigs harboring the fifty longest synteny blocks were compared to themselves using SynMap (https://genomevolution.org/coge/SynMap.pl) and visualized as a dot plot with each syntelog pair shown a dot color-coded based on the synonymous substitution rates (*Ks*) between the pair. The histogram (upper panel) shows both the color keys and distribution for *Ks*. Example synteny blocks representing *P australis*-specific duplication and the ρ duplication shared among the grass lineage are marked with arrows.

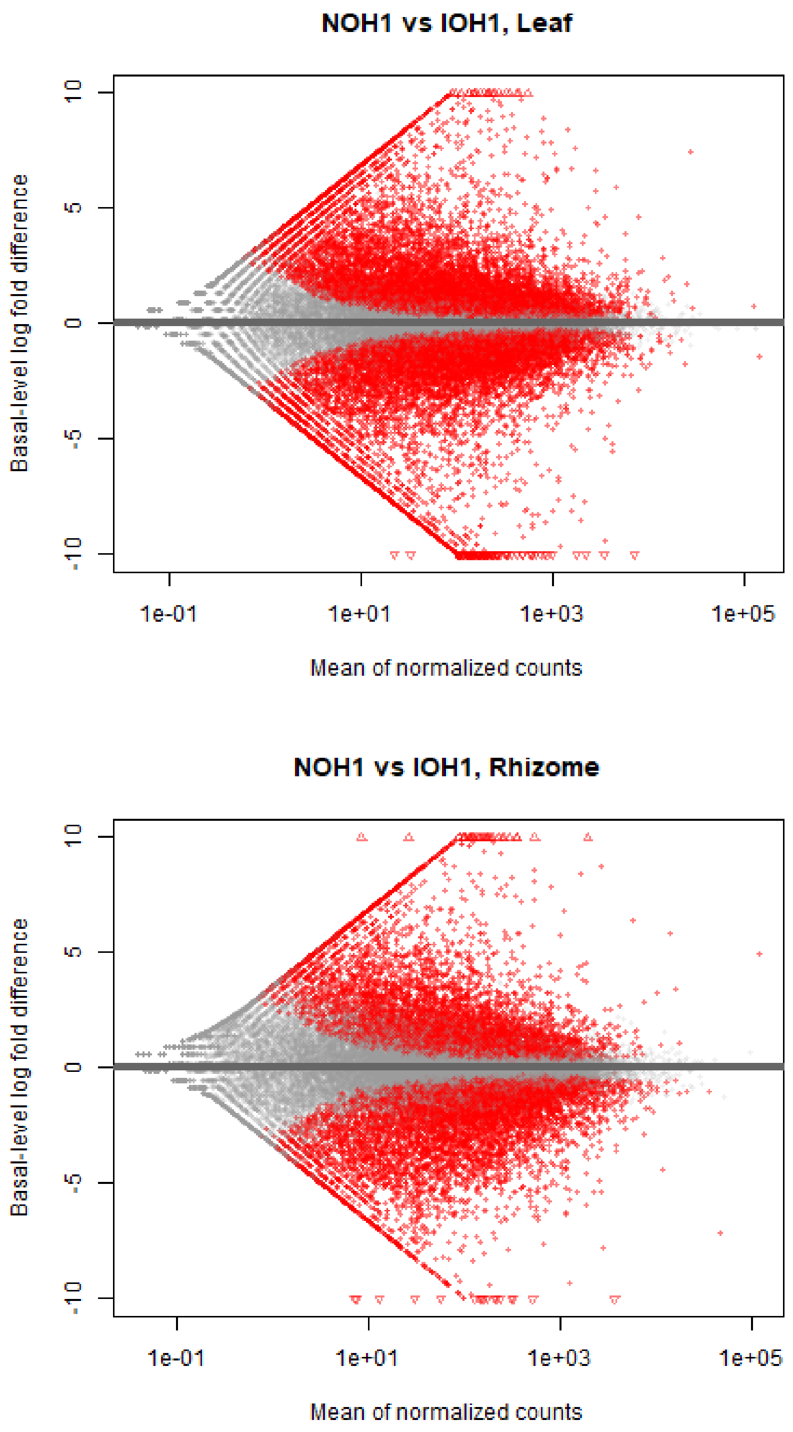


**Supplementary Fig. 4 | RNA-seq analysis comparing basal-level expression among native and invasive *P. australis* genotypes.** Representative "MA"-plots [66] showing distribution of $\log_2$ fold differences (y-axis) and mean normalized RNA-seq counts (x-axis) for NOH1 and IOH1 genotypes. RNA-seq reads from both NOH1 and IOH1 genotypes were aligned to the reference genome sequence and basal-level expression between genotypes were compared after normalization using total number of aligned reads, as detailed in Methods. Reference genes with significantly different basal-level expressions (adjusted p-value<0.05, estimated by DESeq2 [66]) are in red. Note that despite the relatively lower percentage of total reads aligned to the reference genome sequences among native genotypes compared to invasive genotypes (Supplementary Table 4), the MA-plots did not show notable shifts towards lower basal-level expression in the native genotype. All other native and invasive genotype pairs showed similar patterns, corroborating the similar numbers of significantly differentially expressed genes (DEGs) between native and invasive genotype pairs in Fig. 5d.

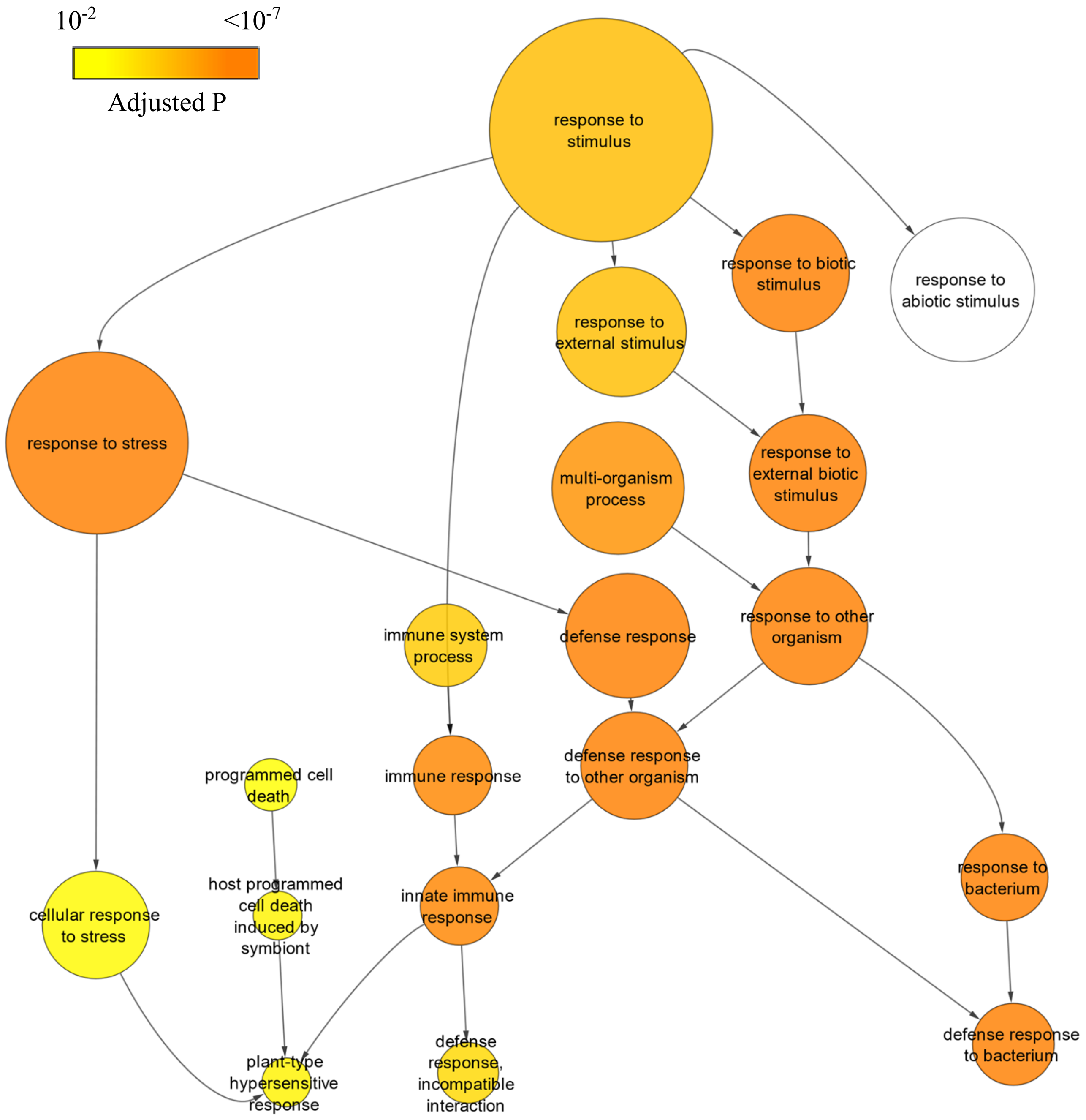


**Supplementary Fig. 5 | An example network of Gene Ontology (GO) terms enriched among *P. australis* reference genes with higher basal-level expression in an invasive genotype.** The network generated using BiNGO (Maere et al., 2005) shows colors indicating the adjusted p-value of enrichment among *P. australis* reference genes with higher basal-level expression in the leaf samples of the IOH1 than the NOH1 genotype. Each GO term was indicated as a circle node, with its diameter mirroring the size of the GO term, and connected to its parent and child GO terms. Note that among the child GO terms of "response to stimulus," defense-related GO terms, e.g. "defense response" (red box), showed the highest levels of enrichment, while other child terms such as "response to abiotic stimulus" was not significantly enriched. For number of genes in each GO terms and detailed p-values for genes showing different basal-level expression between all genotype pairs, see Supplementary Dataset 4.

The following datasets are available at the USGS repository (https://doi.org/10.5066/P91XSF3):

**Supplementary Dataset 1.** *P. australis* version 1.0 reference gene models and syntelog pairs

**Supplementary Dataset 2.** Ortholog groups identified among *P. australis* and five monocots

**Supplementary Dataset 3.** GO clusters enriched among *P. australis* ortholog groups showing conserved or increased gene copy numbers due to *P. australis*-specific duplications

**Supplementary Dataset 4.** RNA-Seq raw read counts and expression rank percentiles for *P. australis* gene models for invasive and native genotypes.

**Supplementary Dataset 5.** RNA-Seq results comparing basal-level expression of *P. australis* genes among invasive and native genotypes

**Supplementary Dataset 6.** GO terms enriched among *P. australis* genes with significantly different basal-level expression among invasive and native genotypes.

**Supplementary Dataset 7.** RNA-Seq results for endophyte-responses of *P. australis* genotypes

**Supplementary Dataset 8.** GO terms enriched among *P. australis* genes whose expression were significantly induced or repressed by endophyte inoculation.
